# Canonical pathoadaptive cystic fibrosis genes in *Pseudomonas aeruginosa* are not CF-specific

**DOI:** 10.64898/2026.08.19.745763

**Authors:** Iris Irby, Elijah C. Mehlferber, Sam P. Brown

**Affiliations:** School of Biological Sciences, Georgia Institute of Technology, Atlanta, Georgia, USA

## Abstract

Research on *Pseudomonas aeruginosa* adaptation in cystic fibrosis (CF) has historically relied on comparing chronic isolates to laboratory reference strains, or evolving reference strains in environments simulating chronic CF. This work has established a small set of genes, including *lasR*, *mucA*, and *mexZ*, as canonical markers of CF pathoadaptation. However, without broad non-CF comparators, it remains unclear how specific these signatures are to CF.

We used a structured literature review to define 20 historically emphasized “canonical CF genes”, then evaluated their mutational patterns across 4,475 genetically distinct *P. aeruginosa* genomes from seven defined clinical and environmental contexts. We tested four competing hypotheses: (1) enrichment in adult CF alone, (2) in adult and pediatric CF combined, (3) in chronic lung infections broadly (including non-CF bronchiectasis), or (4) no strong environment-specific enrichment.

We found little evidence that canonical gene mutations were specifically enriched in adult CF or CF more broadly. Instead, loss-of-function and individual mutations in genes including *mucA*, *mexB*, and *mexZ* were enriched across chronic lung infections, while most canonical genes (including *lasR*) showed no strong environment-specific enrichment.

These results demonstrate that a canon of genes believed to drive pathoadaptation in CF instead largely reflects the narrow comparative framework of past studies rather than CF-exclusive selection. Our findings emphasize shared evolutionary pressures between CF and non-CF bronchiectasis, highlighting opportunities to exchange research and therapeutic insights across chronic infection clinical contexts.

**IMPORTANCE:** *Pseudomonas aeruginosa* infects many sites in the human body, but is studied most intensively in the lungs of people with CF. For decades, researchers compared strains from people with CF against laboratory/reference strains and identified a set of genes that change as the bacterium adapts to the CF lung. These genes have since guided much of the field’s research. Examining thousands of genomes spanning other lung diseases, non-lung human infections, and environmental sources, we show that these changes are not specific to CF. Most historically emphasized CF genes showed no significant CF-specific enrichment, while the clearest enrichment signals mark chronic lung infection broadly, not CF alone. What looked like environment-specific adaptation largely reflects the narrow comparative lens of earlier studies. Encouragingly, insights on *P. aeruginosa* in CF may extend to other chronic lung infections. More broadly, our findings show that claims of niche-specific adaptation require comparison against diverse alternative environments.

## INTRODUCTION

*Pseudomonas aeruginosa* (PA) is an environmental opportunistic pathogen capable of colonizing and persisting across a wide range of environments: from soil and water bodies to industrial and hospital settings, as well as infecting animal and plant hosts. In humans, PA primarily causes infections in immune- or barrier-compromised individuals, colonizing burns, wounds, and lung sites (1). PA infections are of particular clinical concern as they represent a major driver of morbidity and mortality in people with cystic fibrosis (CF) and a frequent cause of chronic infection in people with non-CF bronchiectasis (hereafter bronchiectasis) (2).

Although PA can be isolated from diverse environmental and human-associated contexts, studies of its adaptation in humans have been dominated by work on isolates from chronic CF infections. Most of these studies compare small numbers of chronic CF isolates to reference strains (PAO1, PA14, and others, (3–6)). A growing number of studies take a complementary experimental evolution approach to examine the adaptation of lab strains to CF-mimicking *in vitro* environments (7, 8). Across all these approaches, studies rarely compare strains from different disease stages (e.g., pediatric vs adult CF), non-CF human infections or environmental reservoirs (9, 10). This narrow focus has shaped how the field of PA CF microbiology interprets adaptation: mutations in genes such as *lasR*, *mucA*, and *mutS/L* are often treated as definitive hallmarks of CF pathoadaptation (11–15). However, because robust comparative data are lacking, it remains unclear whether these features are truly distinctive of CF pathoadaptation or potentially more broadly distributed across environments.

Recent work challenges the idea that historically emphasized CF pathoadaptive genes are truly specific to cystic fibrosis (9, 10, 16). In bronchiectasis, expanded genome sampling of PA has shown that strains can harbor genomic features commonly associated with chronic CF infection, suggesting shared evolutionary adaptations across chronic lung environments (17). Broader comparative genomic analyses across thousands of PA isolates also indicate that consistent environment-associated genetic changes appear across diverse human and non-human contexts, including across chronic infections (16). In these studies, mutations in genes such as *mucA* and *mexA/B* appear to be indicative of chronic lung infection more broadly, while other genomic features are identified as distinguishing between chronic infection types (16). These findings suggest that the current canon of “CF pathoadaptive” genes may reflect the narrow comparative frame of prior studies, rather than unique processes of pathoadaptation to the CF lung. These findings raise questions about which genetic signatures are truly CF-specific versus those that reflect broader selective pressures of chronic lung infection, or are simply common across PA regardless of environment. We therefore lack a clear accounting of the genetic signatures underpinning CF-specific pathoadaptation, as compared to those that might be stage-specific, shared across chronic lung infection, or distributed across environmental contexts.

To address the uncertainty surrounding CF-specific versus broader adaptive signatures, we conducted a review of published literature to identify PA genes commonly reported to be associated with CF pathoadaptation. We call these genes ‘canonical CF genes’ to reflect their history of association with CF. We then evaluated the distribution of mutations in these canonical CF genes across diverse infection and non-infection environments. Specifically, we evaluated four hypotheses concerning the range of environments over which these mutations are enriched. Mutations in these genes are either: (1) enriched in adult CF alone, (2) enriched in adult and pediatric CF, (3) enriched across chronic lung infections spanning both CF stages and bronchiectasis, or (4) broadly found across other human and environmental contexts. Clarifying which mutations are unique to CF and which are shared with other human infections is essential for identifying reliable intervention targets and for extending therapies developed for CF-associated infections to other clinical contexts, and vice versa. Beyond our findings in PA, this work establishes a generalizable approach, specifically highlighting the utility of broad environmental contrasts, that can be applied to study environment-specific adaptation across other bacterial species.

## RESULTS

### Defining a canonical CF gene set

To identify the genes most strongly emphasized by the field as markers of CF pathoadaptation, we first conducted a structured literature review of review articles (see methods). We identified 20 genes that were explicitly linked to CF adaptation in at least three of eight retained reviews (Figure 1A; (14, 18–24)). This literature-derived set spans major functional pathways historically associated with CF, including quorum sensing and virulence (*lasR*); mucoidy and biofilm formation (*mucA, algU, algG, pelA*); DNA repair (*mutS, mutL*); antibiotic resistance (*mexZ, mexY, mexX, mexA, mexB, gyrA, gyrB, oprD, nfxB, ampC*); motility (*rpoN*); and metabolism (*aceE, aceF*)

**Figure 1.**
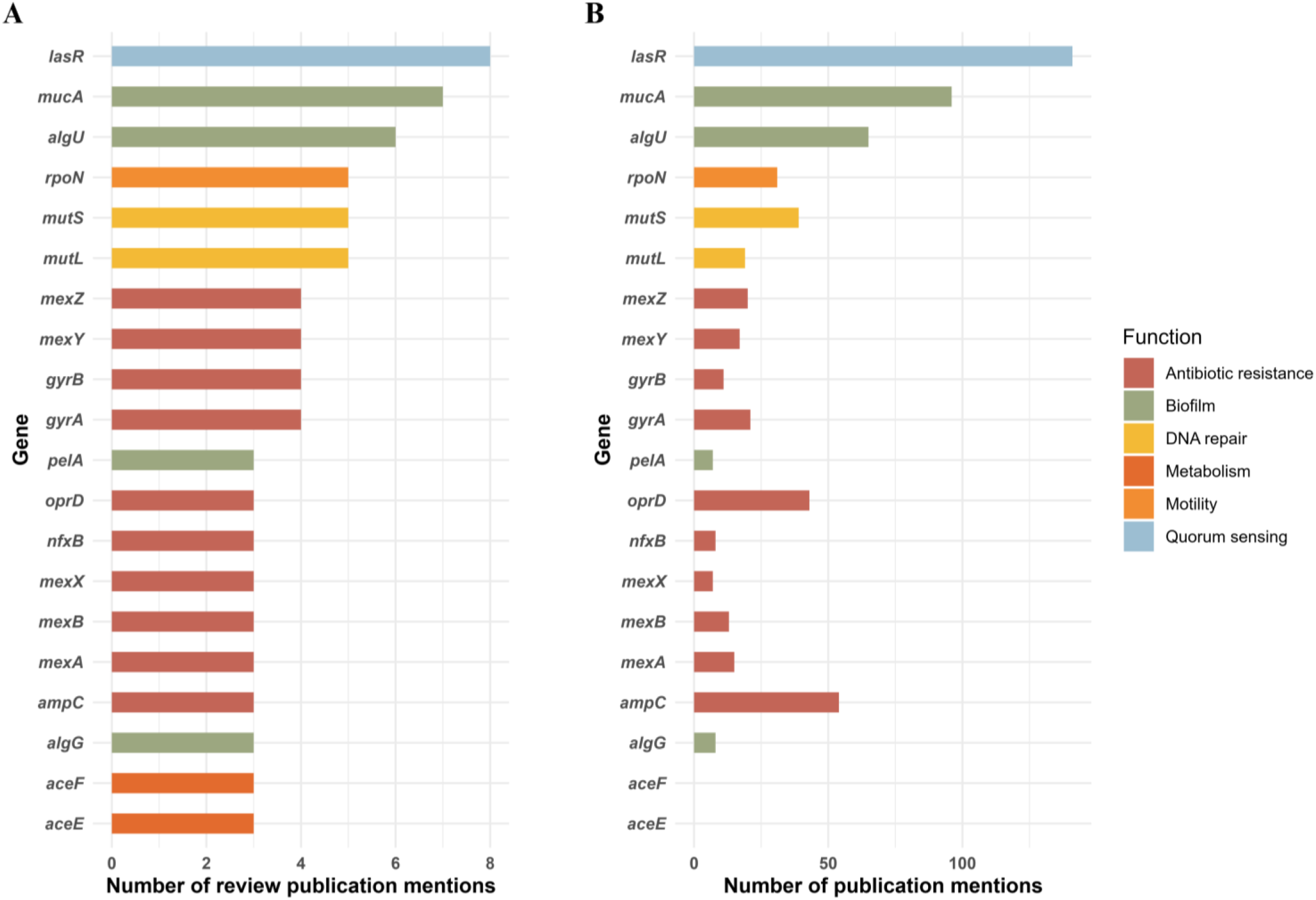
Canonical CF genes span multiple functional categories. (A) Our set of 20 canonical CF genes was defined by a structured literature review (pubmed search string: (((*Pseudomonas aeruginosa*[Title]) AND (cystic fibrosis[Title])) OR (CF[Title])) AND (adapt*[Title]), retaining genes mentioned by gene name in relation to CF pathoadaptation in the article text in at least 3 of the 8 retained review articles. The 20 genes spanned multiple functions, including QS, biofilm, and AMR. Bars show the number of review articles citing each gene in this context (B) Broader literature prominence of the same 20 genes. For each gene, we quantified the number of publications returned by the search string (“Pseudomonas aeruginosa“[MeSH Terms] OR (“Pseudomonas“[All Fields] AND “aeruginosa“[All Fields]) OR “Pseudomonas aeruginosa“[All Fields]) AND (“Cystic Fibrosis“[MeSH Terms] OR “Cystic Fibrosis“[All Fields] OR “CF“[Title/Abstract]) AND “[gene]“[Title/Abstract] where [gene] was substituted with each canonical gene of interest.

We next asked whether this review-derived gene set also captured the broader emphasis of the CF literature. For each of the 20 genes, we quantified its occurrence in publications also addressing PA and CF (Figure 1B; methods). The same genes that dominated the review literature, led by *lasR*, *mucA* and *algU*, were also prominent across the wider literature. Notably, *aceE* and *aceF* are not referenced as the subject of any independent studies, with all review papers citing Marvig et al. 2015 (25). Collectively, the 20 genes accounted for over 600 gene-publication associations, representing 442 unique papers in total (∼140 for *lasR* alone). Thus, Figure 1B broadly validates our review-based definition of the CF canon while also emphasizing how extensively these genes have been studied in a CF context. We therefore used this canonical gene set to ask whether its apparent association with CF persists when compared against isolates from other clinical and environmental contexts.

### Assembling a diverse genomic dataset uncovers clade-level environment structuring

To test our hypotheses, we assembled a database of 4,475 genetically distinct PA genomes with well-defined environmental metadata, expanding upon a previous framework (16). To enable meaningful comparisons while ensuring statistical power, we consolidated environmental source annotations into seven functional categories, distinguishing chronic lung environments (adult CF, pediatric CF, and bronchiectasis) from acute clinical (human infection and pneumonia) and non-clinical (environmental and human-environment) contexts. Because sampling biases and clonal resampling can distort mutational frequencies, we strictly filtered for genome quality and removed clones (defined at 99.99% ANI, (Supplemental Figure 1)).

We hypothesized that phylogenetic population structure could confound the identification of environmentally associated mutations if lineages were unevenly distributed across environments, a factor often overlooked in smaller-scale studies. To provide a clear example of this potential confounder, we evaluated the environmental distribution of the two major PA clades. We found significant differences in environmental membership; clade A was enriched in chronic respiratory environments (adult CF and bronchiectasis), whereas clade B was more associated with acute clinical sources (Figure 2, Supplemental Table 1). Notably, pediatric CF was not significantly enriched across either clade. Because these environmental associations can occur at both broad and fine evolutionary scales, it is critical to account for phylogenetic background across subsequent analyses. We addressed this concern by including high-resolution genomic clusters (genomovar) as a random effect in our gene association models and performing phylogenetic depth analysis on individual mutations as detailed below (26).

**Figure 2.**
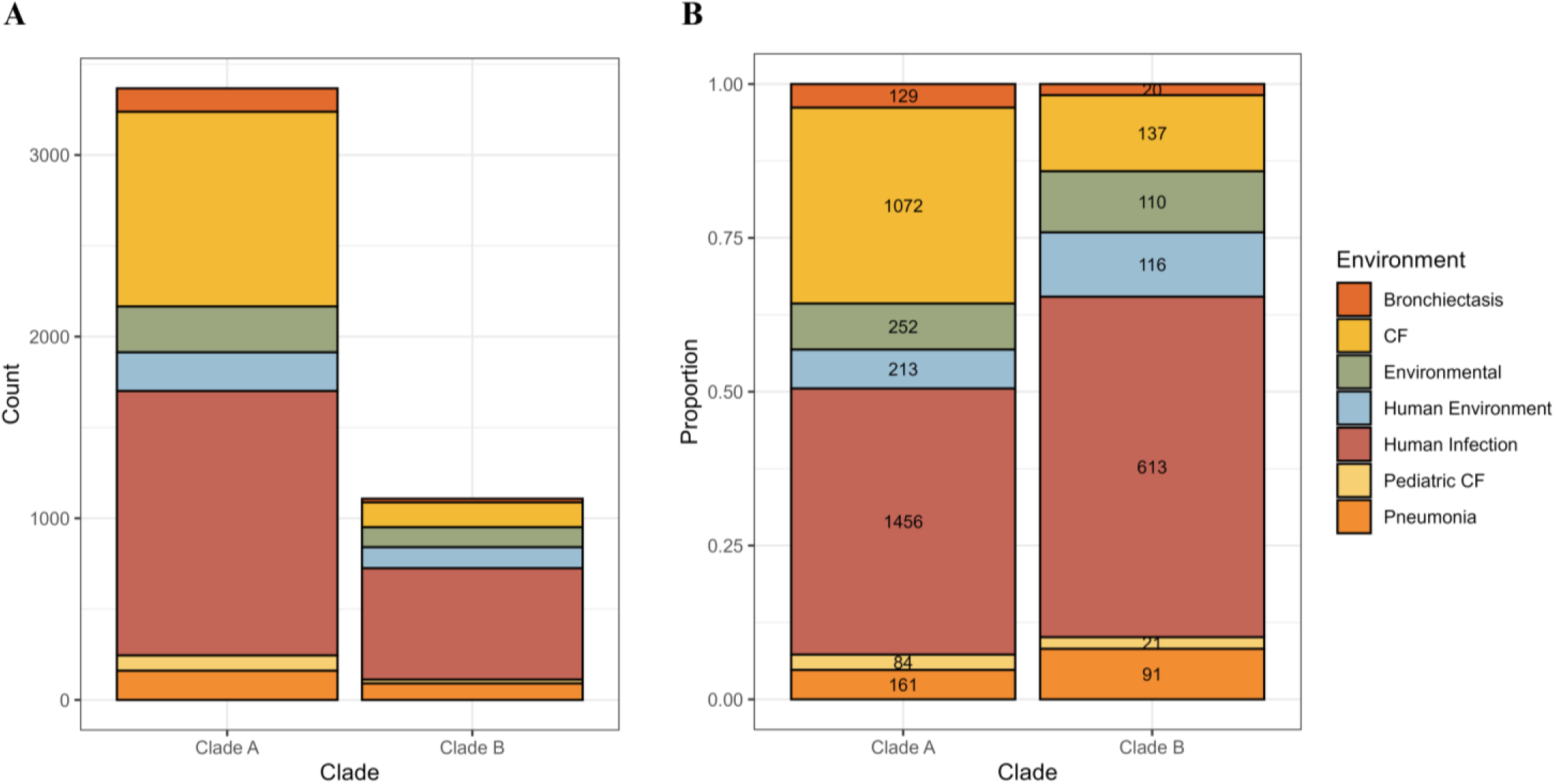
PA genome sequences span diverse environments, with phylogenetic bias in chronic versus acute frequencies. (A) Overall genome counts of PA clades A and B, colored by environment. (B) Environmental proportions of genomes in clades A and B with counts, showing the overall environmental distribution differed significantly between clades (chi-square = 127.8, p < 0.001, Cramer’s V = 0.16). Pairwise Fisher’s exact tests with Bonferroni correction revealed significant clade A enrichment in adult CF (30.3% vs. 10.8%; p < 0.001) and bronchiectasis (5.4% vs. 3.3%; p = 0.034) isolates. Conversely, clade B was significantly enriched in human environment (10.5% vs. 6.3%; p < 0.001), other human infection (55.3% vs. 43.2%; p < 0.001), and pneumonia (8.2% vs. 4.8%; p < 0.001) sources. Genomes were sorted to remove clones (at an ANI threshold of 99.99%) and missing environmental metadata (see methods, Supplemental Figure 1 for data before clonal filtering).

### Loss-of-function profiling reveals a shared chronic lung signature rather than CF-specific pathoadaptation

To test whether mutations in these canonical CF genes are truly specific to CF or shared across other contexts (Hypotheses 1–4), we first profiled mutational enrichment across our seven environmental categories. We prioritized putative loss-of-function (LoF) mutations of the translated gene sequences (including full deletions, frameshifts, and premature stops) as they offer an unambiguous marker of functional differentiation (Supplemental Figure 2). Unsupervised hierarchical and *K*-means clustering of these enrichment profiles revealed a primary, clear separation between chronic lung environments (adult CF, pediatric CF, and bronchiectasis) and all other clinical or environmental contexts (Figure 3). This clustering pattern provided initial support for Hypothesis 3, suggesting that the mutational pattern in canonical CF genes is driven by the broad chronic lung environment rather than CF-specific biology, a trend primarily led by LoF mutations in *mucA, mexB,* and *mexZ*.

**Figure 3.**
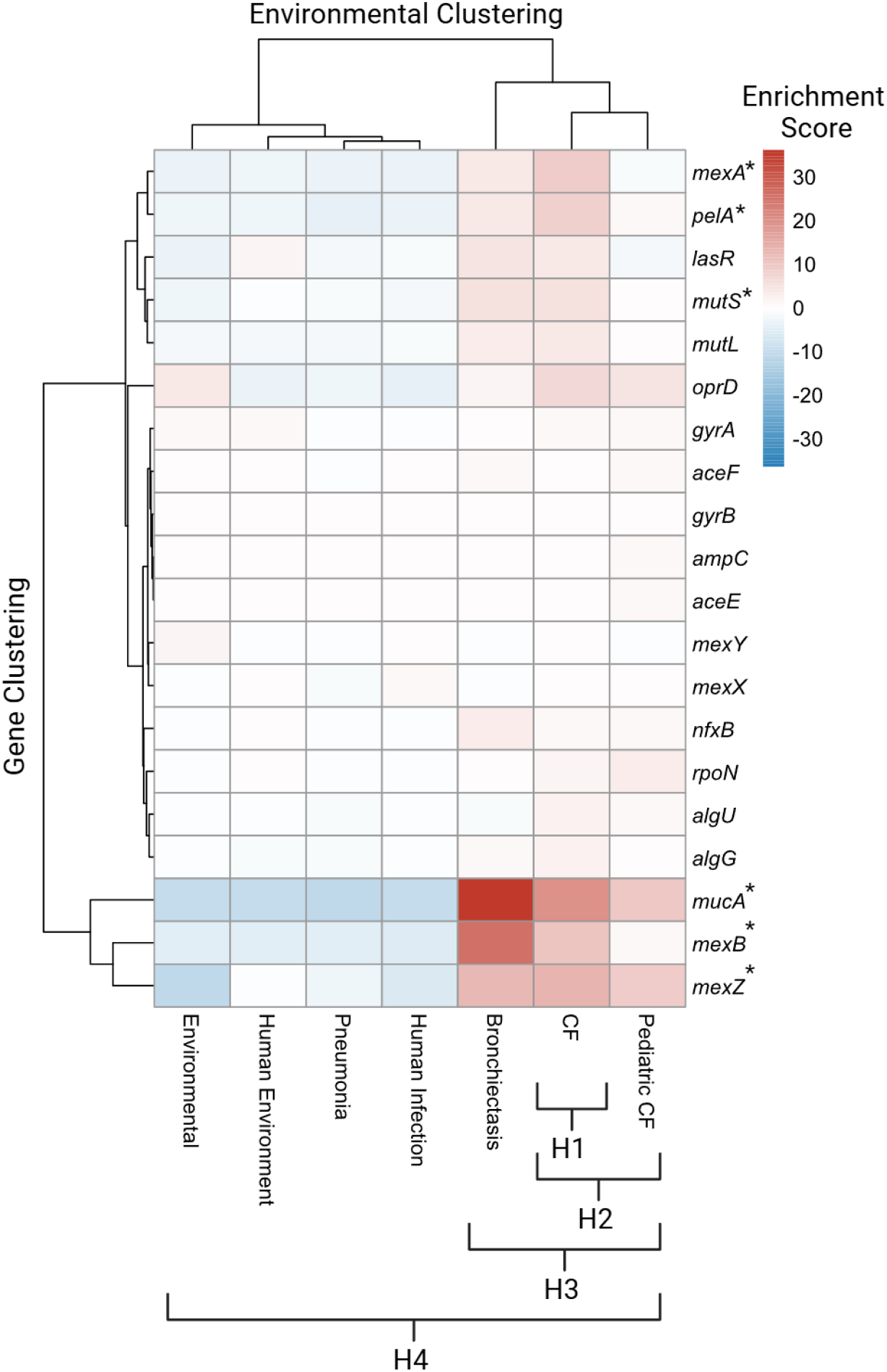
Canonical CF genes are generally not specific to CF. Rows correspond to canonical CF genes, columns correspond to environments. The heat map shows relative mutation enrichment for every gene and environment, where the enrichment score (heatmap) is defined as the difference between the percent of the genomes that are non-functional for a given environment, subtracted from the average percent of non-functional genomes across environments. A higher score (red) indicates enrichment of non-functional mutations in a given gene and environment, while a lower score (blue) indicates depletion of non-functional mutations in that gene and environment. Asterisks mark genes that are identified by the generalized linear mixed effects statistical model as significantly enriched in chronic lung environments (Supplemental Figure 3).

Because broad clustering can be confounded by the lineage-specific environmental biases noted above, we applied gene-level mixed-effects models to formally test our four core hypotheses while accounting for population structure (Supplemental Figure 3). Using a conservative threshold to filter out minor effects (odds ratio ≥ 2, Holm-adjusted *P* < 0.05), we found no evidence that individual genes are uniquely enriched in adult CF (Hypothesis 1) or selectively enriched across combined CF cohorts relative to bronchiectasis (Hypothesis 2). Instead, the statistical models indicated that observed patterns supported a shared chronic lung signature (Hypothesis 3): six genes exhibited significantly elevated LoF mutations and passed our filters across all chronic lung contexts compared to other environments (Figure 3). The strongest enrichment signature was observed for the anti-sigma factor gene *mucA*, followed by the multidrug efflux genes *mexA, mexB,* and *mexZ*, the biofilm associated gene *pelA* and the mismatch repair gene *mutS*. The remaining 14 canonical CF genes either showed no significant environment-specific enrichment of LoF protein variants or failed our filtering thresholds, showing no support for Hypotheses 1, 2, or 3, and indicating that mutations in these genes are found widely across environments.

### Variant-level mutational profiling reinforces a shared chronic lung signature driven by parallel evolution

Because gene-wise LoF patterns showed broad patterns across chronic lung contexts rather than adult CF specificity, we next tracked identical, recurrent amino acid changes across strains. Profiling specific variants potentially provides greater resolution to uncover subtle selective differences within chronic lung environments and highlights precise mutations driving these phenotypes. To isolate robust signals, we evaluated variant significance against fold enrichment, applying a strict false discovery rate threshold (*q* < 0.01), a minimum twofold change (|log_2_(Fold Enrichment)| ≥ 1), and a requirement that the variant appear in at least 10% of genomes within the focal group (Figure 4). We evaluated these thresholds across our primary environmental contrasts (Hypotheses 1–4).

**Figure 4.**
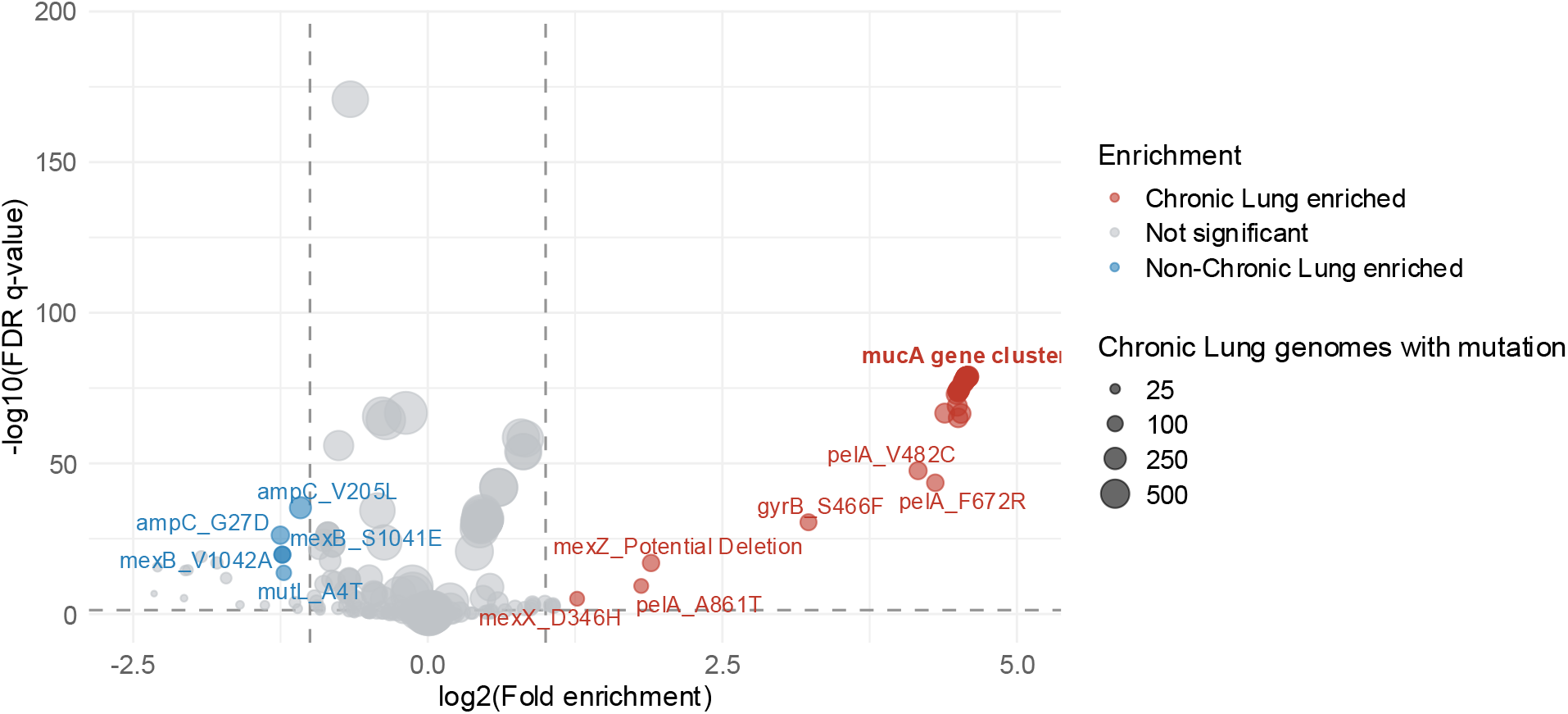
Enrichment of specific mutational variants across chronic lung environments. Specific mutations are plotted by fold enrichment (log2 scale) against statistical significance (-log10 FDR q-value). Horizontal and vertical lines indicate significance and effect-size thresholds (q < 0.01, log2(fold) > +/- 1). Candidate genes are filtered to only those with > 10% prevalence in the chronic group. Each point represents a distinct amino acid mutation, with point size proportional to the variant count within the target environment. Tightly linked, co-occurring variants in MucA are clustered together

Our variant-specific analysis strongly reinforced the shared chronic lung signature while failing to support CF-specific pathoadaptation. Strikingly, no individual mutations were identified as enriched in adult CF vs. other chronic lung, or adult and pediatric CF vs. bronchiectasis comparisons, and the vast majority (49/56) of mutations identified in the adult CF vs. non-chronic or adult CF and pediatric CF vs. non-chronic comparisons were shared by the chronic lung vs. non-chronic group (Supplemental Table 2).

Instead, we saw a suite of individual mutations emerge across all chronic lung environments combined (Hypothesis 3; Figure 4, red points). This focal enrichment was dominated by a dense cluster of MucA variants (likely due to a single frame shift mutation), a recurrent potential deletion in MexZ, and specific individual modifications including PelA (*V*482*C*, *F*672*R*, *A*861*T*), GyrB (*S*466*F*), and MexX (*D*346*H*). Crucially, the relative abundance of these variants remained elevated across non-CF chronic lung contexts, demonstrating that this signature is not merely an artifact of high CF sample representation (Supplemental Table 2). Many proteins identified in our broader gene-level analysis did not yield specific enriched variants, indicating that while the target genes are shared, different strains take distinct mutational routes to achieve modification.

To determine whether these enriched mutations represent deeply rooted, lineage-specific traits or recent evolutionary adaptations, we used ConsenTRAIT (27) to measure their phylogenetic depth across the tree. For all chronic lung-enriched mutations, with the sole exception of MexX (*D*346*H*), the observed phylogenetic depth was statistically indistinguishable from a null expectation generated by randomly shuffling the tree tips (Supplemental Figure 4). This lack of significant clustering indicates that these mutations are distributed broadly and shallowly across the phylogeny. Together, these results suggest that the individual mutations associated with PA strains from chronic lung environments represent independent, parallel mutations occurring across diverse genetic backgrounds, likely in response to the chronic lung environment.

## DISCUSSION

In this study, we identified 20 genes containing mutations that are widely reported to be associated with CF pathoadaptation (Figure 1). We examined the distribution of mutations in these genes across a comprehensive collection of PA strains isolated from diverse environments (Figures 3 and 4), finding little evidence that these mutations are associated with adult CF alone (Hypothesis 1) or adult and pediatric CF (Hypothesis 2). Instead, our results suggest that environment-specific mutation patterns, where they exist, serve as markers of chronic lung infection more broadly (Hypothesis 3), while many of these mutations are not environment-specific and instead are widespread across PA strains regardless of origin (Hypothesis 4). Specifically, LoF mutations in *mucA, mexZ, mexA, mexB,* pelA, and to a lesser extent, *mutS*, were enriched across all chronic lung environments (adult CF, pediatric CF, and bronchiectasis). Both LoF and specific individual mutations displayed substantial overlap among these environments. Together, these results suggest that many mutations commonly described as “CF adaptive” are more accurately interpreted as potential adaptations to chronic lung infection in general.

These findings have important implications for our understanding of bacterial evolution during chronic lung infections. CF and bronchiectasis share core pathological features, including permanent bronchial dilation and sustained inflammatory damage (28), which likely impose similar selective pressures on invading pathogens. The clearest evidence of this shared selection was observed in *mucA*. In all chronic lung environments, we frequently observed *mucA* LoF mutations, which are strongly associated with alginate production and the mucoid phenotype (13). Similarly, shared antibiotic exposures likely drive other dominant phenotypes observed in our study (29, 30). These include resistance mediated by LoF mutations in the *mex* efflux pump genes, and frameshifts in *oprD* associated with carbapenem resistance (31).Our analysis revealed no mutant variants exclusive to the adult CF cohort. Instead, protein variants in MucA, MexZ, PelA, GyrB, and the phylogenetically structured MexX were broadly associated with chronic lung disease (Figure 4), mirroring our LoF analysis (Figure 3).

These data support the conclusions of Mehlferber et al. (16) in two key ways: first, our results also identify a pattern of shared mutations across chronic lung associated genes (specifically MucA and MexZ); second, our results also failed to identify canonical CF genes as distinctive markers (16), while now providing a clear definition of canonical CF genes (Figure 1). While our focus on canonical CF genes precluded examination of non-canonical genes, Mehlferber et al. took a complementary pan-genome approach, showing that multiple non-canonical genes can, in combination, reliably discriminate between adult CF, pediatric CF and non-CF bronchiectasis (16). Among these multi-genic signatures, the strongest markers included *tonB2* (pediatric CF) and *phzD1* (non-CF bronchiectasis). Together, these findings confirm that canonical CF genes are not uniquely tied to CF pathoadaptation.

Beyond supporting a pattern of shared chronic lung pathoadaptation, these results challenge key assumptions regarding adult CF specific pathoadaptation. Many genes frequently cited as hallmarks of adult CF pathoadaptation, most notably *lasR* (32), do not serve as robust diagnostic markers in our dataset. We found little evidence of their enrichment in adult CF isolates specifically, or in chronic lung environments generally, aligning with observations by O’Connor et al. (33). Crucially, this does not preclude their changes from being meaningful in clinical contexts, simply that their occurrence was often comparable to that observed in non-CF environments. This pattern highlights the importance of comparator choice when identifying adaptive mutations. Relying solely on longitudinal CF cohort studies or evolution of model strains in CF-like environments may overestimate CF specificity if mutational patterns also occur across diverse niches. Our findings suggest that broad environmental context should be integrated into future work defining niche-specific adaptation.

Several limitations should be considered when interpreting these results. Our analysis represents a targeted audit of a set of candidate canonical genes, rather than an unbiased genome-wide search for adaptive mutations. Consequently, our conclusions address the specificity of previously proposed CF-adaptive genes, rather than the full spectrum of mutations that may contribute to CF or chronic lung pathoadaptation (see Mehlferber et al. 2026 (16) for an unbiased pangenome survey of local adaptation across CF and non-CF environments). Additionally, because our dataset was aggregated across multiple independent studies, it is enriched for chronic human isolates and contains fewer acute or environmental samples. This aggregation also precluded us from controlling for longitudinal sampling intervals or precise clinical disease state. While this limited metadata resolution may introduce some sources of error, our sample size nonetheless spans a broad range of ecological niches, and the observed mutational patterns remained highly consistent across the evaluated genes.

In summary, this study highlights three key conclusions. First, the canonical CF genes that are enriched in an environment specific manner are enriched broadly across chronic lung infections, suggesting that insights from adult CF pathoadaptation may inform treatment/management for other chronic PA infections such as bronchiectasis. These results emphasize the value of collaboration across CF and non-CF bronchiectasis research to identify shared mechanisms of PA pathoadaptation and improve treatments for chronic lung diseases. Second, most of the genes and mutations previously considered hallmarks of CF pathoadaptation are neither unique to the adult CF environment nor universally common among adult CF isolates overall. This outcome is not surprising given that historical research on PA pathoadaptation has often lacked diverse, non-CF comparator strains. Finally, while these genes may still be valid targets for clinical intervention regardless of their specificity, these findings underscore the need for future functional studies to evaluate candidate genes within a broader, multi-niche evolutionary framework.

## MATERIALS AND METHODS

### Literature review

A structured literature review was conducted to identify relevant PA genes with mutations associated with pathoadaptation to the CF environment. In November 2025, a PubMed title search (Figure 1A) was performed using the search string (((*Pseudomonas aeruginosa*[Title]) AND (cystic fibrosis[Title])) OR (CF[Title])) AND (adapt*[Title]), restricted to review and systematic review publications. This search resulted in 12 review articles, four of which were excluded based on scope: two investigated inter-species bacterial interactions, one was unavailable as a full text publication, and one was dropped because it did not include

### Pseudomonas aeruginosa

The remaining review articles were examined to identify genes with mutations specifically stated to link to CF pathoadaptation. Twenty genes identified in three or more of these publications were retained for further analysis.

To determine whether these candidate genes were broadly represented across the primary literature, a secondary PubMed search was performed in August 2026 (Figure 1B) using MeSH and Title/Abstract terms, with the search string (“Pseudomonas aeruginosa“[MeSH Terms] OR (“Pseudomonas“[All Fields] AND “aeruginosa“[All Fields]) OR “Pseudomonas aeruginosa“[All Fields]) AND (“Cystic Fibrosis“[MeSH Terms] OR “Cystic Fibrosis“[All Fields] OR “CF“[Title/Abstract]) AND “[gene]“[Title/Abstract], where [gene] was substituted with each canonical gene of interest. Restricting “CF” and “[gene]” searches to Title/Abstract fields minimized non-specific or extraneous hits. A full list of PubMed IDs can be found in Supplemental Data 1.

### Database curation

Genomes and metadata were retrieved and processed as previously described (16). Briefly, 30,949 *Pseudomonas aeruginosa* genomes were downloaded from NCBI Datasets (June 6, 2024). Genomes were filtered for quality (N50 > 100 kb, contigs < 300, CheckM completeness >98%, contamination <2%) and clinical isolation context (Data S2, S3), excluding ambiguous “sputum“-only samples. Pediatric CF, adult CF, and bronchiectasis contexts were manually annotated using age (<20 years) and study metadata. This yielded 11,617 high-quality candidate genomes. Genomes were clustered into genomovars (99.5% ANI) and clones (99.99% ANI) using fastANI (v1.3.4, (34)), and re-annotated in a single batch using Bakta (v1.9.3, (35)). Assembly submitter metadata was retained for downstream bias analyses. For this study, we further filtered these genomes to 4,475 based on relevant environmental metadata (excluding animal and non-specific lung samples), and clonal filtering (selecting a single representative clone, at 99.99% ANI per environment).

### Mutation identification

Reference genes and proteins were identified from *P. aeruginosa* PAO1 (NC_002516.2). Each CF-adaptive gene was identified in all PA genomes using nucleotide BLAST (v2.15.0+, (36)) against the reference genes. Genomes found lacking a gene were labelled as deleted and those with insertions were labelled as incomplete. Genes found to be split across multiple contigs were excluded from the analysis.

Gene regions were extracted with bedtools2 getfasta (v2.30.0, (37)) and translated using transeq from EMBOSS (v6.6.0.0, (38)). The resulting protein sequences were compared to the PAO1 reference proteins using a protein BLAST. BLAST query coverage was calculated to be the length of the alignment divided by the reference protein length, multiplied by the percent identity.

Proteins with a query coverage of 100 were classified as wild-type (WT) or “functional”, whereas those with a query coverage of 40 or less were classified as “non-functional”. Proteins with coverage between 40 and 100 were clustered via CD-Hit (v4.8.1, (39)) at 100% sequence length and identity to collapse identical proteins. The resulting protein clusters were aligned to the protein references using clustal-omega (v1.2.4, (40)).

Amino acid substitution mutations, frameshift mutations, premature stop codons, insertions, and deletions were identified from the alignments using Align.IO and Seq.IO modules from BioPython (v1.83, (41)). Proteins containing frameshift mutations and premature stop codons, along with the previously identified deleted and incomplete proteins, were assigned “no function.” Those with amino acid substitutions were assigned as “alternate function.”

### Statistical Analysis

To evaluate whether PA clades are differentially distributed across environments of origin, we analyzed contingency tables cross-tabulating isolate clade membership against niche origin. An overall test of association was performed using Pearson’s chi-square (*χ*^2^) test of independence, with the effect size and strength of association quantified using Cramér’s *V*. To identify which specific clade–niche combinations drove non-random distribution, standardized residuals (*r*) were computed, with absolute values greater than 2 (|*r*| > 2) considered indicative of significant deviation from expected baseline distributions under independence. To test for niche-specific clade enrichment between clade A and clade B, we constructed 2 × 2 contingency tables for each individual niche (focal niche vs. all other niches by clade) and conducted two-tailed Fisher’s exact tests. To control the family-wise error rate across all niche comparisons, raw *p*-values were adjusted using the Bonferroni correction. Comparisons with a Bonferroni-adjusted *p* < 0.05 were considered statistically significant.

We fit generalized linear mixed-effects models (glmer) for each gene with mutation status as the outcome and niche as the predictor (see mutation counts in Supplemental Data 3), using binomial errors and logit link. Specifically, we modelled the distribution of LoF mutations as compared to WT genes + alternative function mutations. To account for population structure, we included random intercepts for Genomovar_Cluster. Models were fitted using the Laplace approximation (nAGQ = 1) and the bobyqa optimizer. Following model fitting, we estimated niche-specific marginal means using the emmeans framework and evaluated five pre-specified contrasts: (i) adult CF versus all other niches, (ii) chronic lung niches combined (adult CF, pediatric CF, and bronchiectasis) versus all other niches, (iii) adult CF versus the other chronic lung niches combined (pediatric CF and bronchiectasis), (iv) adult CF + pediatric CF versus bronchiectasis, and (v) adult CF + pediatric CF versus all other niches. Genes with mutation counts too sparse to support stable model fitting were excluded from contrast estimation, as such cases produce unreliable log-odds estimates regardless of effect direction (Supplemental Figure 3). Excluded genes were categorized based on the nature of the model failure: those where the model failed to converge were designated as “insufficient data”, whereas models that converged but yielded unstable or extreme effect estimates, defined as SE > 10 or | log(OR) | > 7, were categorized as “degenerate fit.” P-values for these contrasts were adjusted using the Holm method to account for multiple comparisons within this predefined family.

To guard against spurious positive findings driven by confounding rather than true within-niche differences, we computed random effect confounding diagnostics for each gene that was identified as significantly associated with a specific contrast, in the most “specific” contrast it was associated (in practice, all chronic CF vs. non-chronic). These diagnostics evaluated whether significant positive contrasts reflected genuine niche-specific elevation or artifacts of Simpson’s paradox (i.e., a niche being concentrated in low-baseline phylogenetic clusters). Specifically, we computed: (1) the Spearman correlation between the proportion of reference-niche isolates per cluster and that cluster’s overall mutation rate (negative values indicate Simpson’s risk), (2) the mean baseline mutation rate of reference-niche-containing clusters minus the grand mean (negative values indicate the reference niche occupies below-average clusters), and (3) the fraction of mixed clusters (containing both reference and non-reference niches) where the reference niche actually has a higher mutation rate (low values indicate the signal is not supported within clusters). Genes were flagged as potentially confounded if any of the following conditions were satisfied: correlation < −0.3, OR baseline bias < −0.05, OR within-cluster support fraction < 0.5.

To evaluate variant-level environmental enrichment, we tracked identical, recurrent amino acid changes across strains to profile specific mutations across our primary environmental contrasts. Genomes were partitioned into target focal environments, encompassing different combinations of chronic lung contexts (adult CF, adult CF + pediatric CF, and adult CF + pediatric CF + bronchiectasis), and non-focal comparator environments (non-chronic contexts or non-focal chronic lung subgroups). For each variant, prevalence was calculated as the proportion of genomes harboring the mutation within the focal (*P*_focal_) and non-focal (*P*_other_) groups. Relative enrichment was quantified by calculating both the prevalence difference (*P*_focal_− *P*_other_) and the log_2_ fold change between groups. Statistical significance was evaluated using False Discovery Rate (FDR) adjusted *q*-values, which were floored at 10^−300^for numerical stability. To isolate robust signals and limit noise from rare variants, mutations were classified as significantly enriched if they met four concurrent criteria: a minimum total occurrence of 100 genomes across all evaluated contexts, an FDR *q* < 0.01, at least a twofold enrichment (log_2_ fold ≥ 1.0), and a requirement that the variant appear in at least 10% of genomes within the focal group. Mutations satisfying these thresholds were designated as focal-enriched (if log_2_ fold > 0) or non-focal-enriched (if log_2_ fold < 0), while all remaining variants were categorized as non-significant.

To determine whether enriched mutations represented deeply rooted, lineage-specific traits or recent evolutionary adaptations, we quantified the phylogenetic depth of environment-of-origin associations using the ConsenTRAIT algorithm (27). For each environmental trait, ConsenTRAIT identified the deepest internal node where at least 90% of descendant tips possessed the trait, calculating the mean depth (*τ*) as the average branch length from that node to all associated tips. Singletons were assigned a depth equal to half the distance to the nearest internal node to ensure comparability across traits. To test whether environmental clustering exceeded random expectations, trait labels were permuted across the phylogeny over 10,000 iterations to build a null distribution of *τ* values against which observed values were evaluated for significance. Finally, to facilitate biological interpretation, mean branch lengths measured in substitutions per site were converted to percent sequence similarity using the transformation: (1 - 2 * *τ_D_*) × 100%).

## ACKNOWLEDGEMENTS

We would like to thank Canan Karakoç, Maria Martignoni, Ryan Lowhorn, Brendan Shrader, and Hrishikesh Patil for discussion and comments on earlier drafts. We thank the National Science Foundation (2209151 to E.C.M., 2406985 and 2321502 to S.P.B., and DGE-2039655 to I.I.), The Army Research Office (W911NF-23-1-0140 to S.P.B.) and the Cystic Fibrosis Foundation (MEHLFE25F0 to E.C.M. and BROWN23I0 to S.P.B.).

## DATA AVAILABILITY

The source code and related data are available on GitHub (https://github.com/GaTechBrownLab/Mutant-Calling).

## Notes

https://github.com/GaTechBrownLab/Mutant-Calling

### Competing Interest Statement

The authors have declared no competing interest.

