## Supplemental Figures for "Canonical pathoadaptive cystic fibrosis genes in *Pseudomonas aeruginosa* are not CF-specific"

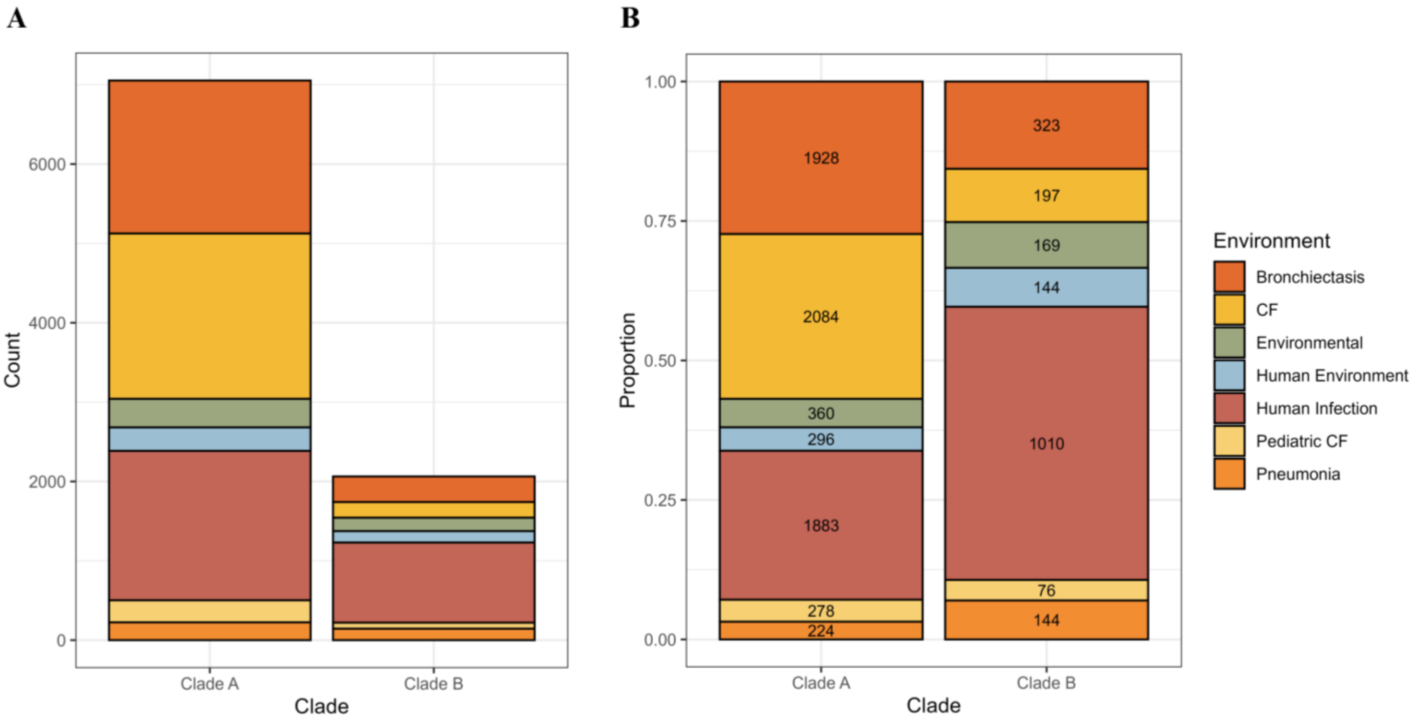


**Supplemental Figure 1. PA genome environmental distribution without dropping clonal genomes.** (A) Overall genome counts of clades A and B, colored by environment. (B) Environmental proportions of genomes in clades A and B with counts, showing the overall environmental distribution differed significantly between clades.

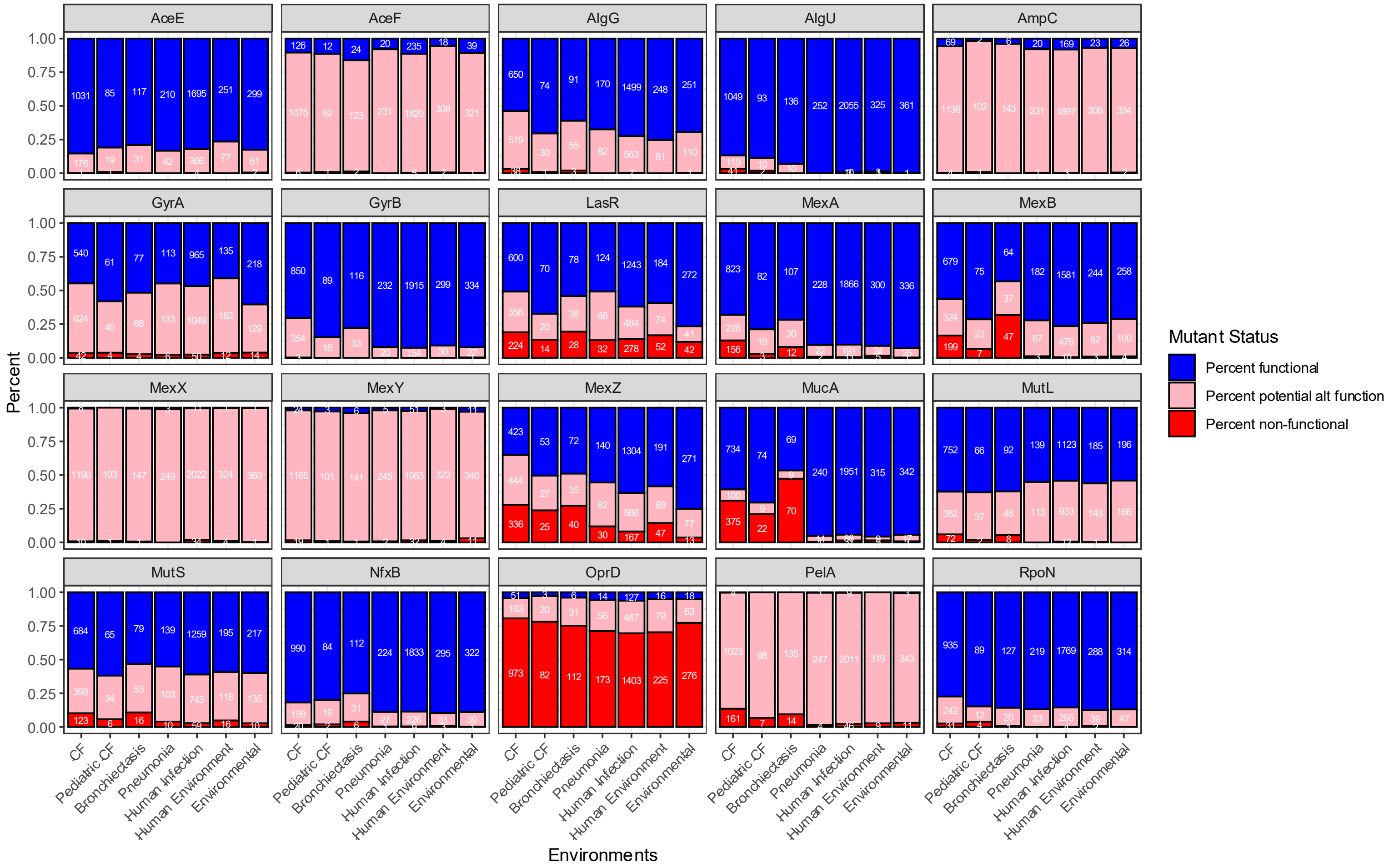


**Supplemental Figure 2. Proportion and count distribution of mutations across proteins and environments.** The proportion of genomes with proteins assigned as functional (blue), alternative function (pink), and non-functional or loss-of-function (red) for every canonical CF gene. Each box corresponds to a canonical CF gene, and each bar corresponds to each environment (CF, pediatric CF, bronchiectasis, pneumonia, human infection, human environment, environmental). Counts for each mutation type per gene and environment are displayed on the bars.

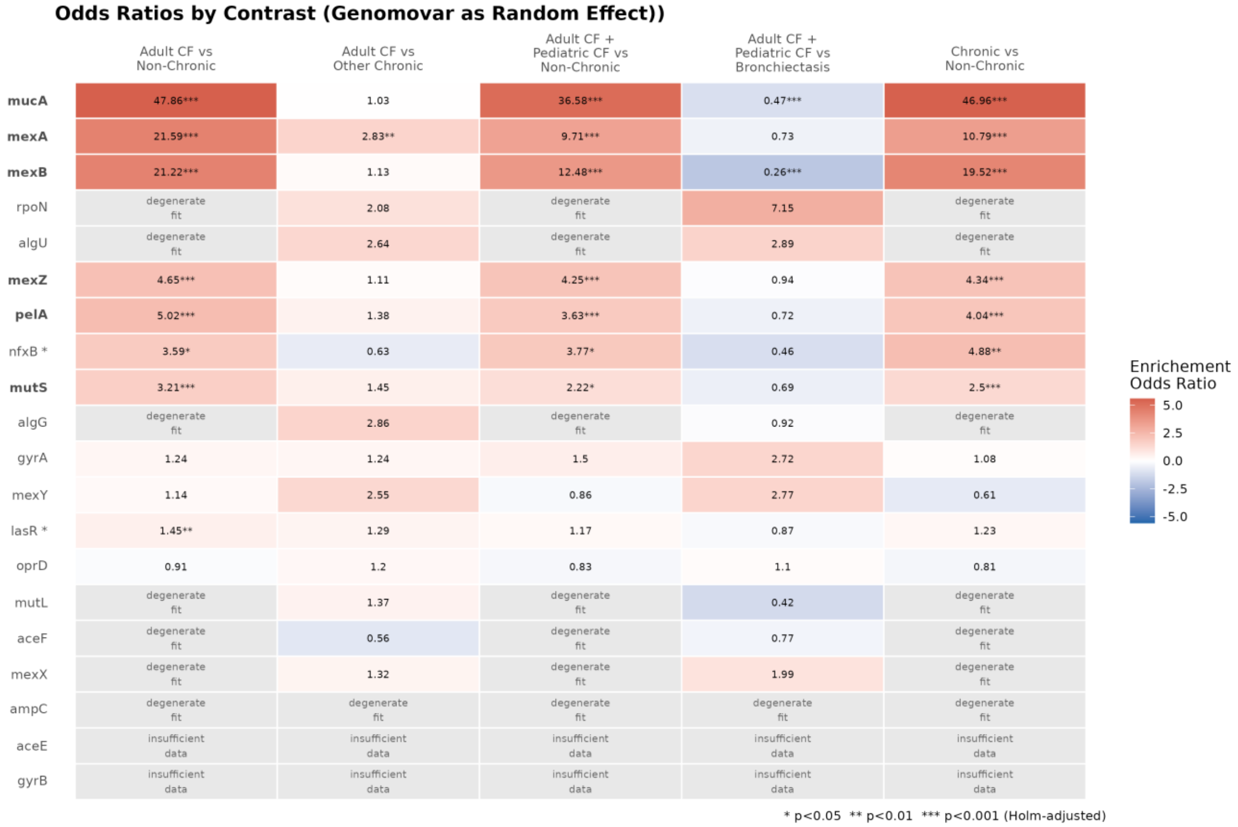


**Supplemental Figure 3. Enrichment of Loss-of-Function (LoF) mutations across environmental contrasts.** Odds ratios from statistical models comparing the distribution of Loss-of-Function (LoF) variants against combined functional and alternative-function variants, colored by log(odds ratio) for symmetry. Each hypothesis was evaluated using distinct focal group comparisons to test for LoF enrichment in adult CF alone (Hypothesis 1; significant enrichment in both adult CF vs non-chronic and adult CF vs other chronic), both adult and pediatric CF (Hypothesis 2; significant enrichment in both adult CF + pediatric CF vs non-chronic and adult CF + pediatric CF vs bronchiectasis), all chronic lung infection cohorts (Hypothesis 3; significant enrichment in chronic vs. non-chronic), or no specific context (Hypothesis 4; no significant enrichment). Meaningful enrichment was defined as an odds ratio >2 with a Holm-adjusted $p < 0.05$, and genes showing significant enrichment were further evaluated for potential confounding factors (e.g., Simpson's paradox, where underlying population structure inflates or distorts effect estimates). After filtering, only Hypothesis 3 is supported by a subset of genes. Genes identified as significant across at least one contrast are bolded, whereas genes that were significant but failed quality filters are marked with an asterisk (*). Specifically, *lasR* was filtered due to a low effect size (OR < 2), while *nfxB* was filtered because its mutations were concentrated in genomovars with low baseline mutation rates, leading the model to inflate its estimated effect (Supplemental Data 2). Genes with too few mutations to support modeling are labeled as either “insufficient data” if the model failed to converge, or “degenerate fit” if the model yielded unreasonable effect estimates (defined as SE > 10 or log(OR) > 7).

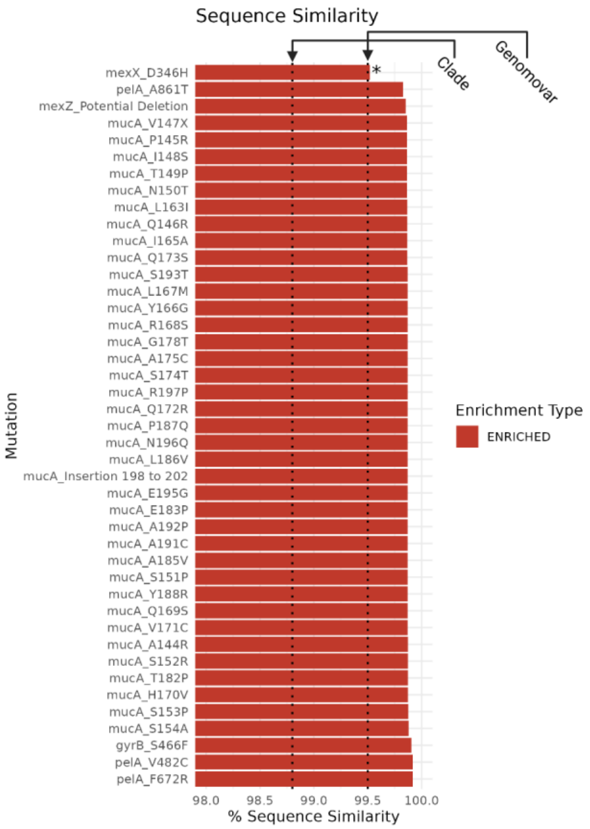


**Supplemental Figure 4. Sequence similarity of strain clusters harboring chronic lung–enriched mutations.** Average sequence similarity of strains within majority single-mutation (>90%) clusters for mutations enriched across chronic lung infections. ConsenTrait average phylogenetic depth was converted to sequence similarity, with reference lines marking the genomovar (99.5%) and clade (98.8%)

thresholds. The vast majority of mutations cluster exclusively below the genomovar level, exhibiting a distribution indistinguishable from random permutation. Only *mexX* D346H forms a significantly deeper cluster than expected by chance.

| Niche | Total_Count | CladeA_Count | CladeB_Count | CladeA_Prop | CladeB_Prop | Std_Residual_A | Std_Residual_B | Fisher_P_Value | Fisher_P_Bonf | Significant |
|---|---|---|---|---|---|---|---|---|---|---|
| Adult CF | 1139 | 1019 | 120 | 0.3026 | 0.1083 | 5.534 | -9.648 | 0.0000 | 0.0000 | ** |
| Bronchiectasis | 219 | 182 | 37 | 0.0541 | 0.0334 | 1.342 | -2.339 | 0.0049 | 0.0343 | ** |
| Environmental | 362 | 252 | 110 | 0.0748 | 0.0993 | -1.234 | 2.152 | 0.0110 | 0.0772 | |
| Human Environment | 329 | 213 | 116 | 0.0633 | 0.1047 | -2.195 | 3.827 | 0.0000 | 0.0001 | ** |
| Other Clinical | 2069 | 1456 | 613 | 0.4324 | 0.5532 | -2.553 | 4.450 | 0.0000 | 0.0000 | ** |
| Pediatric CF | 105 | 84 | 21 | 0.0249 | 0.0190 | 0.562 | -0.980 | 0.3029 | 2.1204 | |
| Pneumonia | 252 | 161 | 91 | 0.0478 | 0.0821 | -2.077 | 3.621 | 0.0000 | 0.0002 | ** |

**Supplemental Table 1. Distribution and differential enrichment of *Pseudomonas aeruginosa* clades across environmental niches.** Summary of isolate counts, within-clade proportions, and standardized residuals across environments of isolation for strains from Clades A and B. Both raw ($p$) and Bonferroni-corrected ($p_{\text{adj}}$) $p$-values are shown; asterisks denote statistically significant enrichment ($p_{\text{adj}} < 0.05$).

| Count | Gene | Mutation | Adult CF vs Non-Chronic | Adult CF vs Other Chronic | Adult CF + Pediatric CF vs Non-Chronic | Adult CF + Pediatric CF vs Bronchiectasis | Chronic vs Non-Chronic |
|---|---|---|---|---|---|---|---|
| 1 | ampC | ampC_G27D | 0.37 (p=8.5e-27) | NA | 0.4 (p=9.2e-26) | NA | 0.42 (p=6.7e-27) |
| 2 | ampC | ampC_V205L | 0.42 (p=4.1e-38) | NA | 0.43 (p=6.5e-39) | NA | 0.47 (p=5.2e-36) |
| 3 | gyrB | gyrB_H148N | 2.14 (p=2.9e-04) | NA | 2.05 (p=7.3e-04) | NA | NA |
| 4 | gyrB | gyrB_S466F | 9.31 (p=4.9e-27) | NA | 8.62 (p=2.6e-25) | NA | 9.39 (p=3.5e-31) |
| 5 | mexB | mexB_S1041E | 0.41 (p=1.0e-17) | NA | 0.42 (p=2.5e-18) | NA | 0.43 (p=1.9e-20) |
| 6 | mexB | mexB_V1042A | 0.41 (p=1.0e-17) | NA | 0.42 (p=2.5e-18) | NA | 0.43 (p=1.9e-20) |
| 7 | mexX | mexX_A30T | 0.5 (p=2.8e-21) | NA | NA | NA | NA |
| 8 | mexX | mexX_D346H | 2.69 (p=1.2e-06) | NA | 2.61 (p=1.4e-06) | NA | 2.41 (p=8.5e-06) |
| 9 | mexY | mexY_N1036T | 0.48 (p=2.9e-15) | NA | NA | NA | NA |
| 10 | mexZ | mexZ_Potential Deletion | 3.63 (p=1.5e-14) | NA | 3.72 (p=5.2e-16) | NA | 3.72 (p=1.1e-17) |
| 11 | mucA | mucA_A144R | 22.18 (p=4.6e-64) | NA | 21.39 (p=6.1e-64) | NA | 22.42 (p=1.1e-73) |
| 12 | mucA | mucA_A175C | 23.95 (p=1.3e-70) | NA | 22.87 (p=1.3e-69) | NA | 23.68 (p=6.1e-79) |
| 13 | mucA | mucA_A185V | 24.09 (p=4.7e-71) | NA | 23 (p=4.9e-70) | NA | 23.79 (p=2.6e-79) |
| 14 | mucA | mucA_A191C | 24.09 (p=4.7e-71) | NA | 23 (p=4.9e-70) | NA | 23.79 (p=2.6e-79) |
| 15 | mucA | mucA_A192P | 24.09 (p=4.7e-71) | NA | 23 (p=4.9e-70) | NA | 23.79 (p=2.6e-79) |
| 16 | mucA | mucA_E183P | 24.09 (p=4.7e-71) | NA | 23 (p=4.9e-70) | NA | 23.79 (p=2.6e-79) |
| 17 | mucA | mucA_E195G | 24.09 (p=4.7e-71) | NA | 23 (p=4.9e-70) | NA | 23.79 (p=2.6e-79) |
| 18 | mucA | mucA_G178T | 23.95 (p=1.3e-70) | NA | 22.87 (p=1.3e-69) | NA | 23.68 (p=6.1e-79) |
| 19 | mucA | mucA_H170V | 23.95 (p=1.3e-70) | NA | 22.87 (p=1.3e-69) | NA | 23.68 (p=6.1e-79) |
| 20 | mucA | mucA_I148S | 22.92 (p=8.1e-67) | NA | 21.79 (p=1.6e-65) | NA | 22.65 (p=1.2e-74) |
| 21 | mucA | mucA_I165A | 23.65 (p=1.6e-69) | NA | 22.6 (p=1.4e-68) | NA | 23.45 (p=5.4e-78) |
| 22 | mucA | mucA_Insertion 198 to 202 | 24.09 (p=4.7e-71) | NA | 23 (p=4.9e-70) | NA | 23.79 (p=2.6e-79) |
| 23 | mucA | mucA_L163I | 23.21 (p=7.2e-68) | NA | 22.06 (p=1.7e-66) | NA | 22.88 (p=1.4e-75) |
| 24 | mucA | mucA_L167M | 24.09 (p=4.7e-71) | NA | 23 (p=4.9e-70) | NA | 23.79 (p=2.6e-79) |
| 25 | mucA | mucA_L186V | 24.09 (p=4.7e-71) | NA | 23 (p=4.9e-70) | NA | 23.79 (p=2.6e-79) |
| 26 | mucA | mucA_N150T | 23.07 (p=2.5e-67) | NA | 21.93 (p=5.2e-66) | NA | 22.76 (p=4.2e-75) |
| 27 | mucA | mucA_N196Q | 24.09 (p=4.7e-71) | NA | 23 (p=4.9e-70) | NA | 23.79 (p=2.6e-79) |
| 28 | mucA | mucA_P145R | 22.92 (p=8.1e-67) | NA | 21.79 (p=1.6e-65) | NA | 22.65 (p=1.2e-74) |
| 29 | mucA | mucA_P187Q | 24.09 (p=4.7e-71) | NA | 23 (p=4.9e-70) | NA | 23.79 (p=2.6e-79) |
| 30 | mucA | mucA_Q146R | 23.65 (p=1.6e-69) | NA | 22.6 (p=1.4e-68) | NA | 23.45 (p=5.4e-78) |
| 31 | mucA | mucA_Q169S | 24.24 (p=4.6e-71) | NA | 23.14 (p=4.9e-70) | NA | 23.9 (p=2.6e-79) |
| 32 | mucA | mucA_Q172R | 24.09 (p=4.7e-71) | NA | 23 (p=4.9e-70) | NA | 23.79 (p=2.6e-79) |
| 33 | mucA | mucA_Q173S | 24.39 (p=2.1e-71) | NA | 23.27 (p=2.5e-70) | NA | 24.02 (p=1.6e-79) |
| 34 | mucA | mucA_R168S | 23.95 (p=1.3e-70) | NA | 22.87 (p=1.3e-69) | NA | 23.68 (p=6.1e-79) |
| 35 | mucA | mucA_R197P | 24.09 (p=4.7e-71) | NA | 23 (p=4.9e-70) | NA | 23.79 (p=2.6e-79) |
| 36 | mucA | mucA_S151P | 21.16 (p=3.2e-60) | NA | 20.18 (p=3.0e-59) | NA | 20.93 (p=2.1e-67) |
| 37 | mucA | mucA_S152R | 22.81 (p=1.4e-58) | NA | 21.79 (p=8.3e-58) | NA | 22.65 (p=7.4e-66) |
| 38 | mucA | mucA_S153P | 23.14 (p=1.1e-59) | NA | 22.09 (p=7.5e-59) | NA | 23.03 (p=2.7e-67) |
| 39 | mucA | mucA_S154A | 22.4 (p=5.0e-61) | NA | 21.36 (p=4.4e-60) | NA | 22.53 (p=9.7e-70) |
| 40 | mucA | mucA_S174T | 24.09 (p=4.7e-71) | NA | 23 (p=4.9e-70) | NA | 23.79 (p=2.6e-79) |
| 41 | mucA | mucA_S193T | 23.95 (p=1.3e-70) | NA | 22.87 (p=1.3e-69) | NA | 23.68 (p=6.1e-79) |
| 42 | mucA | mucA_T149P | 22.92 (p=8.1e-67) | NA | 21.93 (p=5.2e-66) | NA | 22.76 (p=4.2e-75) |
| 43 | mucA | mucA_T182P | 23.51 (p=5.7e-69) | NA | 22.46 (p=4.5e-68) | NA | 23.33 (p=1.6e-77) |
| 44 | mucA | mucA_V147X | 22.92 (p=8.1e-67) | NA | 21.79 (p=1.6e-65) | NA | 22.65 (p=1.2e-74) |
| 45 | mucA | mucA_V171C | 24.39 (p=2.1e-71) | NA | 23.27 (p=2.5e-70) | NA | 24.02 (p=1.6e-79) |
| 46 | mucA | mucA_Y166G | 23.95 (p=1.3e-70) | NA | 22.87 (p=1.3e-69) | NA | 23.68 (p=6.1e-79) |
| 47 | mucA | mucA_Y188R | 24.24 (p=4.6e-71) | NA | 23.14 (p=4.9e-70) | NA | 23.9 (p=2.6e-79) |
| 48 | mutL | mutL_A4T | 0.38 (p=2.7e-14) | NA | 0.4 (p=5.0e-14) | NA | 0.43 (p=2.1e-14) |
| 49 | oprD | oprD_E185Q | 0.5 (p=5.2e-30) | NA | NA | NA | NA |
| 50 | oprD | oprD_G425A | 0.47 (p=1.9e-12) | NA | 0.5 (p=1.2e-11) | NA | NA |
| 51 | oprD | oprD_P186G | 0.5 (p=5.2e-30) | NA | NA | NA | NA |
| 52 | oprD | oprD_V189T | 0.5 (p=6.7e-30) | NA | NA | NA | NA |
| 53 | pelA | pelA_A861T | 3.73 (p=7.3e-10) | NA | 3.58 (p=1.8e-09) | NA | 3.51 (p=4.9e-10) |
| 54 | pelA | pelA_F672R | 21.56 (p=1.2e-43) | NA | 20.49 (p=9.9e-43) | NA | 19.8 (p=2.7e-44) |
| 55 | pelA | pelA_V446I | 2.13 (p=7.0e-05) | NA | 2.05 (p=1.4e-04) | NA | NA |
| 56 | pelA | pelA_V482C | 19.34 (p=3.8e-47) | NA | 18.31 (p=6.0e-46) | NA | 17.89 (p=2.5e-48) |

**Supplemental Table 2. Enrichment of individual mutations across environmental contrasts.** Similar to the gene-level Loss-of-Function (LoF) analysis, individual mutation enrichment was evaluated across focal group comparisons to test for enrichment in Adult CF alone (Hypothesis 1), Adult and Pediatric CF combined (Hypothesis 2), all chronic lung infection cohorts (Hypothesis 3), or no specific context (Hypothesis 4). The resulting odds ratios, and their associated Bonferroni-corrected ($p_{\text{adj}}$) $p$-values are shown. Only mutations identified as significantly enriched across at least one comparison are shown. Although some mutations were enriched in CF relative to non-chronic controls, they lacked significant differential enrichment when compared to other chronic environments, failing the criterion for CF-specific adaptation (Hypotheses 1 and 2). Instead, the vast majority of enriched mutations occurred across all chronic lung groupings (supporting Hypothesis 3), consistent with convergent chronic adaptation.
